# Terrestrial 3D Laser Scanning for Ecosystem and Fire Effects Monitoring

**DOI:** 10.1101/2024.04.09.587551

**Authors:** Mary Carlton Murphy, E. Louise Loudermilk, Scott Pokswinski, Brett Williams, Emily Link, Laila Lienesch, Leta Douglas, Nicholas Skowronski, Michael Gallagher, Aaron Maxwell, Grant Snitker, Christie Hawley, Derek Wallace, Irenee Payne, Tim Yurkiewicz, Andrew J. Sánchez Meador, Chad Anderson, J. Mark Jackson, Russell Parsons, Melissa Floca, Isaac Nealey, Ilkay Altintas, J. Kevin Hiers, Jon Wallace

## Abstract

Long-term terrestrial ecosystem monitoring is a critical component of documenting outcomes of land management actions, assessing progress towards management objectives, and guiding realistic long-term ecological goals, all through repeated observation and measurement. Traditional monitoring methods have evolved for specific applications in forestry, ecology, and fire and fuels management. While successful monitoring programs have clear goals, trained expertise, and rigorous sampling protocols, new advances in technology and data management can help overcome the most common pitfalls in data quality and repeatability. This paper presents Terrestrial Laser Scanning (TLS), a specific form of LiDAR (Light Detection and Ranging), as an emerging sampling method that can complement and enhance existing monitoring methods. TLS captures in high resolution the 3D structure of a terrestrial ecosystem (forest, grassland, etc.), and is increasingly efficient and affordable (<$30,000). Integrating TLS into ecosystem monitoring can standardize data collection, improve efficiency, and reduce bias and error. Streamlined data processing pipelines can rigorously analyze TLS data and incorporate constant improvements to inform management decisions and planning. The approach described in this paper utilizes portable, push-button TLS equipment that, when calibrated with initial transect sampling, captures detailed forestry, fuels, and ecological features in less than 5 minutes per plot. We also introduce an interagency automated processing pipeline and dashboard viewer for instant, user-friendly analysis, and data retrieval of hundreds of metrics. Forest metrics and inventories produced with these methods offer effective decision-support data for managers to quantify landscape-scale conditions and respond with efficient action. This protocol further supports interagency compatibility for efficient natural resource monitoring across jurisdictional boundaries with uniform data, language, methods, and data analysis. With continued improvement of scanner capabilities and affordability, these data will shape the future of terrestrial ecosystem monitoring as an important means to address the increasingly fast pace of ecological change facing natural resource managers.

## INTRODUCTION

Monitoring, as the “intermittent recording of the condition of a feature of interest to detect or measure compliance with a predetermined standard,” is an often-mandated framework for the agencies tasked with stewarding our public lands and natural resources (Hellawell 1991). Monitoring is essential to ecosystem management (Christensen and others 1996) and for informing management goals under accelerating climate change (Hiers and others 2012). In this paper, we describe the application of emerging technology to improve the utility and efficiency of monitoring within terrestrial ecosystems (forests, grasslands, etc.). Under the Forest and Rangeland Renewable Resources Planning Act of 1974, later amended by the National Forest Management Act of 1976, Congress formally directed the U.S. Department of Agriculture Forest Service to monitor and assess the Nation’s natural resources (Morrison and Marcot 1995). At the national scale, the Forest Inventory and Analysis (FIA), Forest Health Monitoring (FHM), and FEAT/FIREMON Integrated (FFI) programs conduct surveys or utilize remote sensing data to assess the status and trends of various forest and other vegetation (grasslands) conditions. Similarly, the Department of the Interior supports monitoring through the Bureau of Land Management Assessment, Inventory, and Monitoring (AIM), and National Park Service and U.S. Fish and Wildlife Service Inventory & Monitoring (I&M) programs. Localized monitoring programs vary considerably by discipline, but can have multiple, concurrent sampling protocols for wildlife, forestry, and wildland fuels management programs. Programs such as these play a crucial role in adaptive resource management by evaluating management outcomes and the effects of natural disturbances to measure trends and departures from a predetermined desired state (Legg and Nagy 2006). Monitoring can ensure accountability by tracking the condition of valuable resources such as timber and endangered species habitat. It also documents the effects of land management activities, including timber harvesting, fuel treatments, and controlled burns. Effective monitoring data communicates management actions and goals and can be harnessed for public education as well as institutional or agency support.

Successful monitoring programs are goal-oriented, have rigorous sampling protocols, require trained expertise, and have a diligent remeasurement schedule (Lindenmayer and Likens 2010, Vos and others 2000). Furthermore, they require innovation to address the present and future challenges facing natural resource management, such as climate change and urbanization (Lovett and others 2007). Intensive, comprehensive, and detailed sampling protocols promote precision and accuracy, while minimizing user bias and measurement error, all creating high-quality datasets that are useful for informing management and agency decisions as well as research and development. Challenges with monitoring that can contribute to poor data quality are associated with inconsistency in measurements among users, between plots, and through time, all increasing bias and error (Lindenmayer and Likens 2010, Pokswinski and others 2021). This is not withstanding potential changes in sampling protocols through time or across sites, which can reduce repeatability and compatibility between seemingly similar datasets. These issues can occur for various reasons, including but not limited to insufficient resources or time, personnel changes within a given monitoring program, and lack of trained personnel overall or with critical skill sets.

Depending on the monitoring program’s objectives, sampling methods may include distance measures (Bitterlich 1952, Lindsey and others 1958, Picard and Bar-Hen 2007), line intercepts (Brown 1974, Canfield 1941), or quadrat methods (Lindsey and others 1958, Pound and Clements 1898), and standardized forest inventory protocols, such as fixed or variable radius plots. Each of these methods is tailored towards variables of interest. For example, Brown’s (1974) transects, an inventory method for sampling downed, dead woody material, are commonly used for fire effects and fuels monitoring, but do not capture information on live or aerial fuels, herbaceous cover, and canopy structure, and may require high sampling intensities to reduce uncertainty (Sikkink and Keane 2008). Another example, the composite burn index, assigns a severity index based on assumed pre-fire conditions and visual estimates of fuel consumption, greenness, necrosis, and char following burning (Gallagher and others 2021). These sampling approaches provide quick general estimates of variables of interest and may benefit by incorporating more robust quantitative measurements using advanced technology.

Advances in sampling tools and statistics can address many of these monitoring challenges through whole plot remote sensing measurements (Loudermilk and others 2009, Rowell and others 2016, Skowronski and others 2007), and artificial intelligence and machine learning-driven analyses to reduce personnel bias (Ditria and others 2022, McClure and others 2020, Skowronski and others 2011). Terrestrial laser scanning is one such tool to improve and enhance monitoring with output data that is not variable specific, but rather captures the entire structure of a plot and measures individual components (tree boles, shrubs, etc.) with high precision.

### Terrestrial Laser Scanning for Monitoring

Light Detection and Ranging (LiDAR) is an active remote sensing tool used for recording discrete spatial positions in 3D. Numerous types of LiDAR instruments are operated via mobile (MLS), terrestrial (TLS), airborne (ALS), and unmanned aerial systems (UAS), among others. In each system, the scanners collect data by emitting lasers, which bounce off features and return to a sensor 10 to 100s of thousands of times per second (Figure 1), similar to sonar, but at a different electromagnetic wavelength. Scanners use two main techniques for calculating range measurements: phase-shift (or waveform) and time-of-flight (Liang and others 2016, Newnham and others 2015). Phase-shift scanners use a continuous laser to measure the intensity of reflected objects. In contrast, time-of-flight scanners record the time between laser pulses to determine distance with the speed of light. The resulting output of both techniques is a point cloud of data points displaying the 3D environment in x, y, and z coordinate space (Figure 2). Global Positioning Systems (GPS) can associate the point cloud with real Earth coordinates relative to a map projection, and an inertial measurement unit (IMU) can allow for data collection from a moving platform. Many LiDAR systems are also equipped with cameras that capture RGB (true-color) images to complement the 3D point cloud data. As a result, LiDAR technology is an increasingly relevant decision-support tool within the natural resource community (Hudak and others 2009). For example, ALS data produces wall-to-wall digital terrain models (DTM), canopy height models (CHM), and locates, detects, and measures individual or close groups of trees across large landscapes or watersheds.

**Figure 1.**
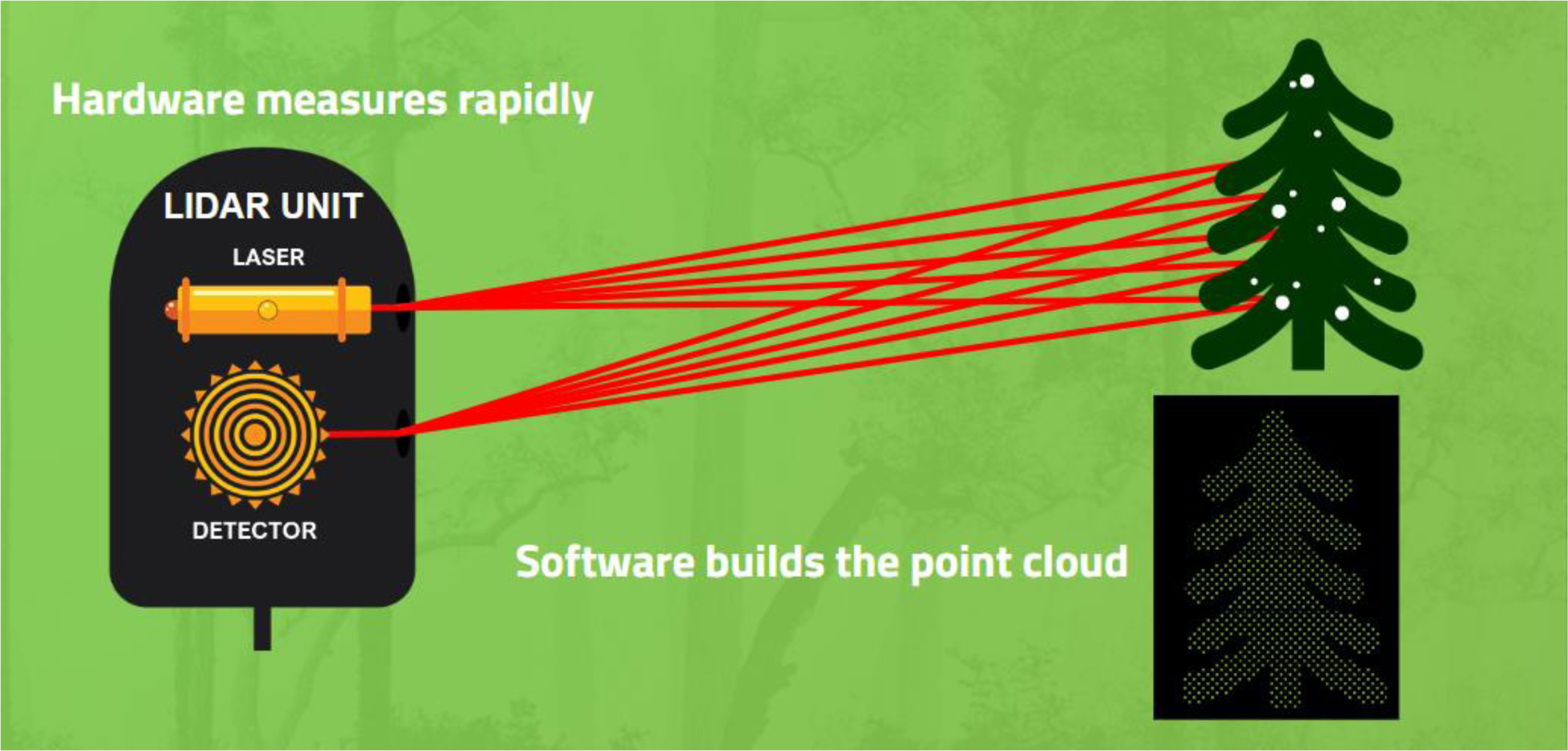
A terrestrial laser scanning (TLS) device sends out rapid laser pulses that bounce off objects and returns to the device’s detector, where distance is calculated using the speed of light. A point cloud image containing thousands of laser points is the output. (Courtesy: Jess Stetson).

**Figure 2.**
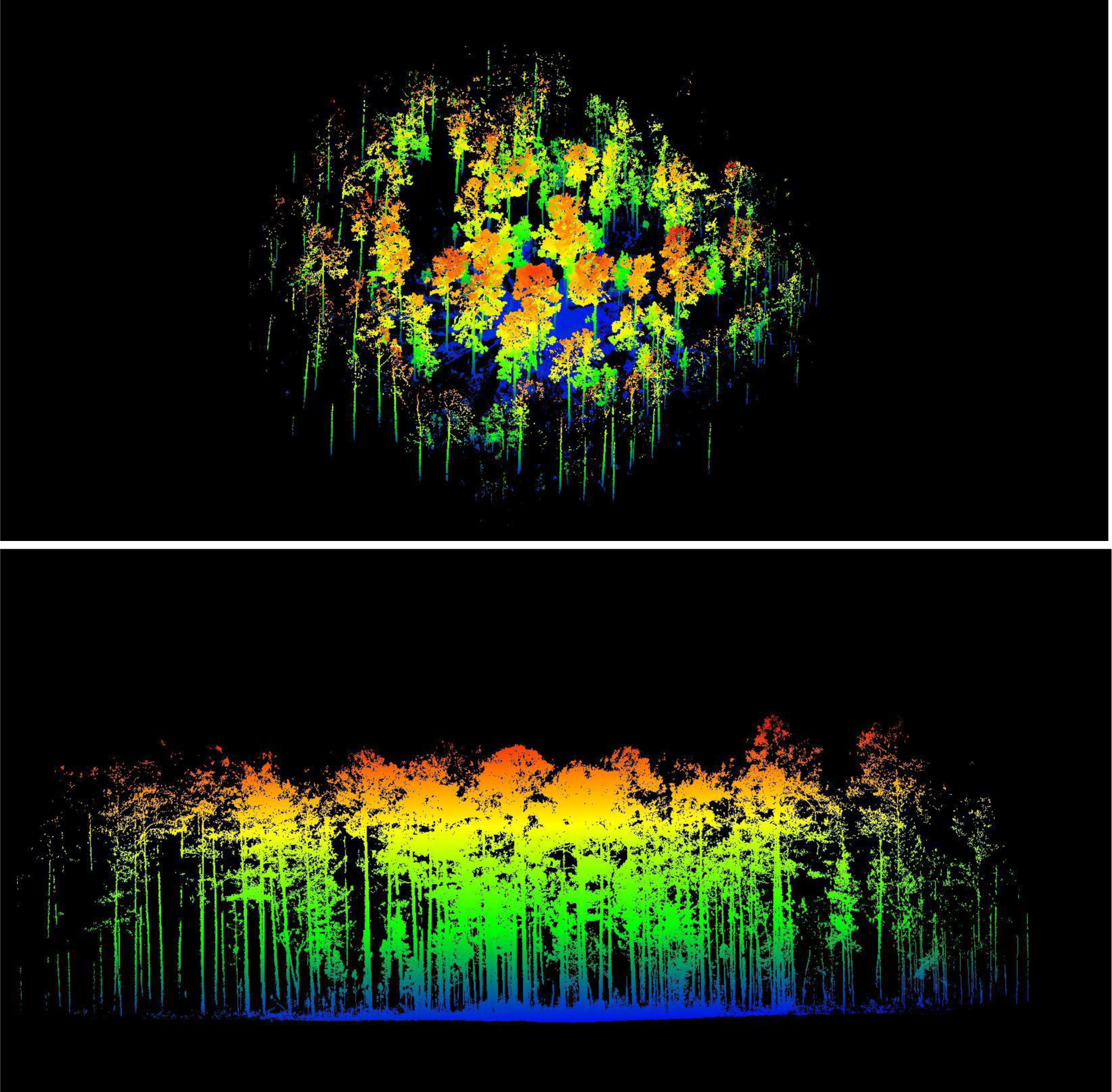
Raw 3D point cloud image output from a single 360° TLS scan of a southern pineland; from birds-eye view (above) and side view (below). For scale, the trees are ∼25m tall. (Courtesy: E. Louise Loudermilk, USFS)

Of these various new approaches, TLS in particular has gained traction for wide natural resource applications. TLS-based monitoring is uniquely poised to standardize data collection with repeatable, quantitative data collection methods and with instruments that are generally lightweight, portable, increasingly affordable, and easy to operate with minimal training (push-button in many cases) (Gallagher and others 2021, Pokswinski and others 2021). This allows for rapid sampling (<5-minutes per scan) of the 3D structure of a typical forest plot (<0.1 ha) down to a centimeter resolution and hundreds of direct output metrics critical to habitat and fuel monitoring such as basal area and canopy cover. Furthermore, these metrics can be used to predict non-measured attributes, such as surface fuel loading and fire severity (see Recent Research). As a result, TLS can monitor large areas while producing a breadth of metrics, allowing for integrated monitoring across sites and disciplines. Repeated site visits are easily conducted with TLS monitoring, making it ideal for adaptive management and supplementing established monitoring plots with revisits. Additionally, 3D point clouds and RGB images derived from TLS instruments offer new visual tools for communicating management treatment effects or plot-level changes that occur through time.

There is a continuous need for rapid and accurate understanding of ecosystem dynamics. Amidst habitat loss, severe weather, increasing wildfire frequency, and climate change, landscapes are changing quickly, challenging adaptive management. This innovative TLS-based monitoring approach advances information for management by optimizing resource efficiency and effectiveness.

### Objectives

This paper introduces a novel TLS monitoring protocol for advanced ecosystem monitoring, associated with the Interagency Ecosystem LiDAR Monitoring (IntELiMon) program.^1^ This approach leverages tripod-based, push-button TLS technology to provide a fast, accurate, and consistent method for quantifying 3D plot structure and providing hundreds of forest and vegetation structural metrics. We provide a guide on the protocol’s technical background, field data collection, and processing options. We also highlight a state-of-the-art web portal for automated analysis and streamlined, interagency-compatible data storage and visualization. We summarize current and future research on the technology, including modeling techniques to monitor fuels and ecosystems, as well as linkages to fire modeling for land management decision support.

## FIELD DATA COLLECTION METHODS

### TLS Instrument

For this monitoring protocol, a stationary TLS instrument is used to capture detailed (centimeter resolution) plot-level data with little to no instrument configuration and easy operation. Nonstationary options (mobile scanners on backpacks, vehicles, or UAS) can cover larger areas and are useful with trained expertise but are difficult to replicate consistently and efficiently. This monitoring protocol using a stationary TLS, includes an automated analysis and processing pipeline, which presently supports data from the Leica BLK360 G1 and G2 (Leica Geosystems Inc., Heerbrugg, Canton St. Gallen, Switzerland). Many TLS instruments are available, however, and research and development into monitoring with other scanners (e.g. FARO and Trimble products) is currently underway (See Recent Research and Applications). Considerations for instrument choice include instrument price, weight, portability, ease of use, field durability, data accessibility, laser specifications, and processing requirements.

All details in this paper pertain to use of the Leica BLK360 G1 (hereafter referred to as “BLK360 G1”) and processing and analyzing its outputs. The BLK360 G1 is a terrestrial laser system that is lightweight (1 kg), affordable (<$30,000), and splash resistant. The unit has an area of coverage that is 360° horizontally and 300° vertically, a range of accuracy of 4-mm at 10-m distance and 7-mm at 20-m distance, and a maximum range distance of 60 m (Table 1, Figure 2). The unit has multiple sampling density settings to control the number of points collected, which can help manage data collection speed and file size (Loudermilk and others 2023). For this monitoring protocol, the medium-density setting is required for compatibility with the automated analysis pipeline, which results in a scan time of less than 5 minutes.

**Table 1.** Specifications and performance of the Leica BLK360 G1 terrestrial laser scanner.

| LEICA BLK360 | SPECIFICATIONS |
| --- | --- |
| Maximum Range | 60m |
| Optimal Plot Range | 10m |
| Accuracy | 6mm @ 10m |
| Point Capture Speed | 360,000 points/s |
| Size | 16cm tall x 10cm diameter |
| Weight | 1kg |
| Scan Time | <5 minutes |

<sup>1</sup> https://dmsdata.cr.usgs.gov/lidar-monitoring/

The BLK360 G1 records only first returns of the laser, meaning that if a laser pulse intersects and bounces off multiple surfaces (split laser beam) to result in multiple laser returns, only the first one is kept. This is important when generating information about the density of porous materials, like leafy forest vegetation, because it allows for automated comparison of expected to actual laser returns and predicts where the density of objects is concentrated. These analyses are useful for predicting everything from fuel density to forestry metrics.

The BLK360 G1 is mounted on a camera tripod and operated by pressing a single button in the field. The simplicity of operation allows for efficient, repeated data collection with minimal training. In less than 5 minutes, the laser scans and collects a dense 3D point cloud (360,000 points per second) with a usable scan radius of 10–15 m. This monitoring protocol sets a ‘usable’ scan radius because of occlusion; while the scanner can capture data up to a 60-m distance, objects inherent to a forested environment (e.g., tree boles, foliage, debris) can occlude or block the scanner from capturing objects behind them, resulting in a shorter distance in which quality data can be captured. Experience and research show that a 10-to 15-m radius is ideal for plot monitoring purposes (Gallagher and others 2021, Loudermilk and others 2023, Pokswinski and others 2021). Despite this ‘cropping’ of the point-cloud, the output provides a significant amount of data. For example, one 10-m radius scan can result in a structural image with approximately 6 to 8 million 3D points (including RGB camera data) within an area similar to a one-tenth acre forestry plot (approximately 0.03 ha or 0.07 acre).

The BLK360 G1 battery life typically lasts for one day of scanning and exporting of data and can easily be supplemented with a battery change. Scan storage capacity is 32 GB or roughly 100 medium-resolution scans. The instrument is light enough to be carried by hand or in a backpack and operates within a temperature range of 5 °C to 40 °C (41 °F to 104 °F) as specified by the manufacturer. Wind and rain can affect scanner operability, potentially damaging the instrument as well as impacting point cloud accuracy. It is thus important to consider temperature and weather extremes during field sampling that may impact scanner functioning and plan accordingly.

### BLK360 G1 Applications

The BLK Data Manager Utility and Leica Cyclone Register 360 PLUS (BLK Edition) (Leica Geosystems Inc., Heerbrugg, Canton St. Gallen, Switzerland) are required for data offloading and file conversion for the BLK360 G1. These requirement guidelines may change over time. Please check with the company and instrument specifications and manuals. Refer to the latest Methods Manual for a list of suggested applications to complement TLS monitoring.^2^

#### A Note: Digital data entry or paper datasheets?

Within the IntELiMon protocol, there is an option to use ArcGIS Survey123 for digital data entry through a tablet^2^ or to use paper datasheets. Ultimately, the user can determine which option works best for their monitoring team and program. ArcGIS Survey123 provides automatic, offline digital datasheet entry that syncs with ArcGIS Online once connected to Wi-Fi. This method requires a field-durable mobile tablet and additional training for formatting and customization. Alternatively, 1-page paper datasheets are handwritten, customizable, do not require additional hardware or technical formatting, yet require manual data entry after field sampling. If mobile tablets are the method of choice, it is suggested to always keep paper datasheets on hand during field sampling in case of technological mishaps or in environmental conditions that may limit mobile tablet use in the field.

### BLK360 G1 Hardware Preparation

Prior to field sampling, check the BLK360 G1 for firmware updates and ensure the stand-alone/push-button functionality is engaged under the Capture Settings. Refer to the latest Methods Manual for instructions on updating the BLK360 G1 firmware, BLK License Management, and setting the BLK360 G1 push-button behavior.^2^

### Field sampling protocol

A tiered sampling protocol has been established for efficient and accurate TLS data collection that is customizable for monitoring needs (Figure 3). Following the protocol steps in order will ensure that scans are not visually obscured by the operator or equipment and that groundcover, tree structure, and fuels are not impacted by trampling during plot setup. Refer to the latest Methods Manual for a sample equipment list for TLS fuels and the sampling of ecological conditions.^2^

**Figure 3.**
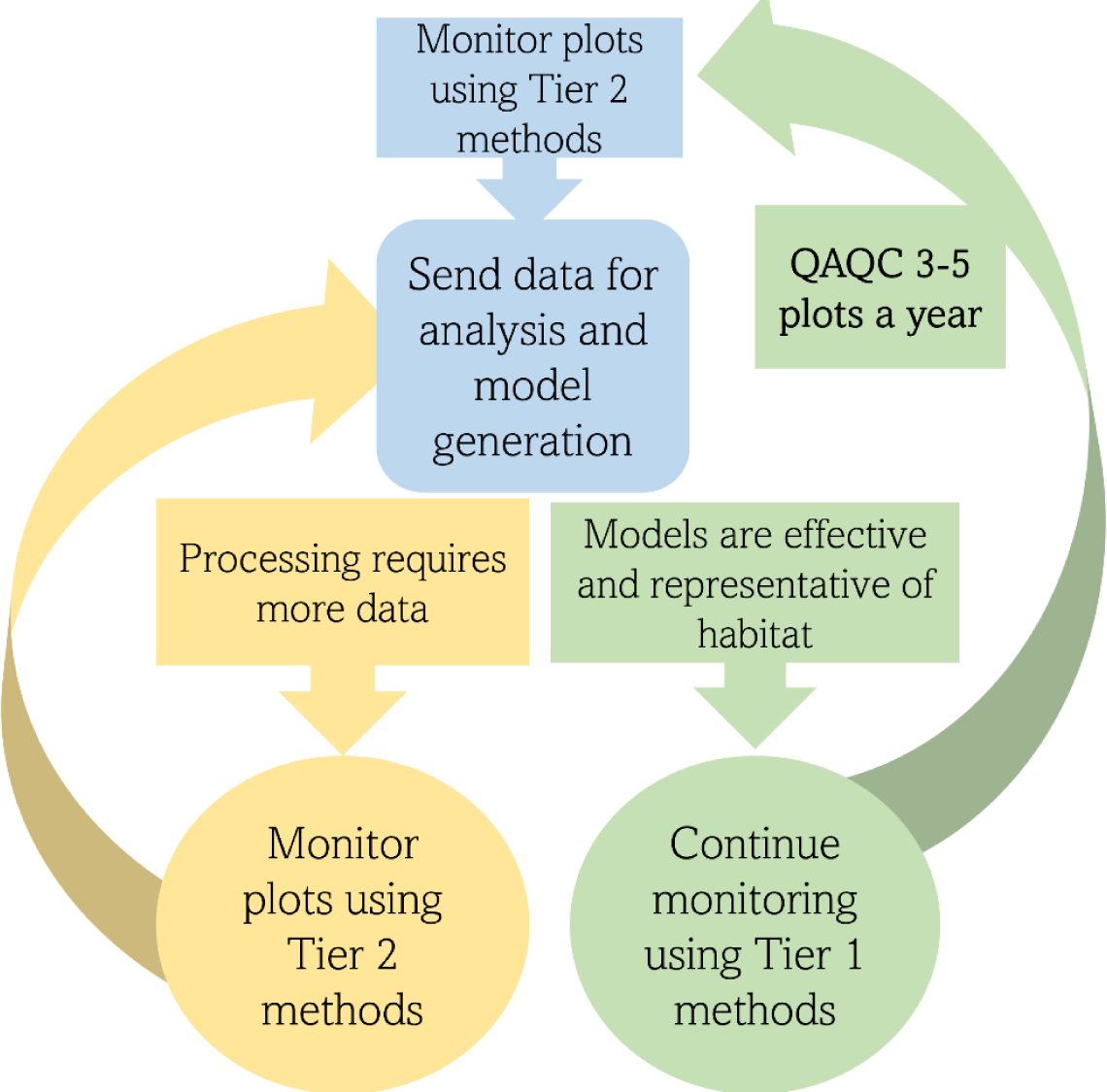
Conceptual diagram of the tiered TLS Monitoring Protocol. QAQC: Quality Assurance Quality Control. (Courtesy: Emily Link, USFWS).

There are three tiers of data collection. **Tier 1** sampling occurs on all plots and includes a single 360° TLS scan and variable-radius sampling using a prism (hereafter referred to as prism pass) for hardwoods and conifers at designated plots. **Tier 2** sampling includes Tier 1 plus additional (and often initial or intermittent) transect fuel sampling and overstory tree identification within each plot. **Tier 3** includes Tier 2 plus any additional desired sampling, often for more site-specific, objective-driven, or research and development needs (see Recent Research and Applications for examples). It is important to note that the goal of Tier 2 sampling is to develop calibration datasets for specific monitoring sites, ensuring that models can accurately predict structural and surface fuel metrics with linear regression analyses. As such, areas that are first implementing TLS monitoring typically require, at a minimum, Tier 2 sampling at as many plots as possible and, following model calibrations, are then able to step down to the simpler Tier 1 in subsequent sampling at the same plots, or often within the same ecosystem with similar structural characteristics.

Once plots are established and the monitoring team is ready, it is suggested to perform trial runs of the sampling protocol. This can be done by creating mock plots in easily accessible areas. Here, the team can run through the protocol, work out the details of plot navigation, set up, and execution for both Tier 1 and Tier 2 sampling. This also allows for practice to minimize trampling, refine fuel categories (if needed), and run through the download and processing steps back in the office. This ensures a comfortable and efficient working environment and high-quality data is collected in the established monitoring plots.

### Plot selection

Plot locations are chosen based on site-specific monitoring objectives, such as a goal to represent ecological variation, forest density, or fuel density conditions across a site. The number of plots can vary highly among sites, dependent on the range of conditions, access, size, burn history and frequency, and available resources. It is important to note that greater numbers of established plots contribute to better models built from Tier 2 sampling as well as more consistent long-term data from Tier 1 sampling. Random, grid, or stratified random plot sampling designs are commonly used at monitoring sites and consulting with a sampling expert can improve monitoring outcomes. Using existing monitoring plots is also suggested, from which legacy data sets may be linked to the TLS data for incorporating past trends. If no monitoring program exists, or resources are low, one can start with a simple plot design (e.g. pick one ecosystem type of interest and assign a manageable number of randomly distributed plots). More plots can be added over time as resources become available and objectives are refined.

Although the scanning range of the BLK360 G1 is extensive and dependent on vegetation density, we suggest using a 10-m radius (314.16 m² or 0.03 ha or 0.078 acre) plot size to coincide with a 10-m radius transect sampling, to minimize occlusion within the single scan, and to maximize detected ground points for height stratification. However, a larger plot size may be used depending on local conditions (e.g., 15-m radius in open forest conditions). Given a 10-m radius plot size, plots should be at least 15-m apart and at least 15-m from water, roads, or other built features. Frequent foot or mechanical traffic and safety concerns (e.g. tree snags) must be considered when selecting plots and throughout the continuous sampling of plots through time. Trampling or other soil and vegetation disturbance will impact the quality and consistency of data collected and models developed over time.

### TLS Target Placard

The BLK360 G1 has no integrated GPS or compass, so 3D data points are not georeferenced or oriented. A GPS unit can be used to determine plot center, but not orientation. If orientation is desired, an optional target placard can be created and placed 2-m outside the plot radius (e.g., 10-m plot radius and the placard at 12-m due north) to orient each scan in cardinal space. Instructions for target placard construction can be found in the Methods Manual.^2^

### TLS data collection with the BLK360 G1

#### Tier 1 Sampling

Navigate to plot center using Field Maps or another mapping tool. If the designated plot center is close to a tree, adjust plot center so that the laser is not within an estimated distance of three times the diameter of the tree, if possible. Adjust plot location for any safety concerns. Record the site location, surveyor initials, date and time, and GPS the location of plot center.

At plot center, a tripod is raised to the maximum height (approximately 1.5-m) and situated so the BLK360 G1 is directly over plot center. The 2-m pole with the attached TLS target placard is firmly secured into the ground 12-m north of plot center (e.g., 2-m outside the 10-m plot radius) (Figure 4). Before collecting TLS scans and any additional physical data, prevention of trampling and disturbance of groundcover, tree structure, and fuels should be considered. For further instructions on scanning with the BLK360 G1, refer to the latest Methods Manual.^2^

**Figure 4.**
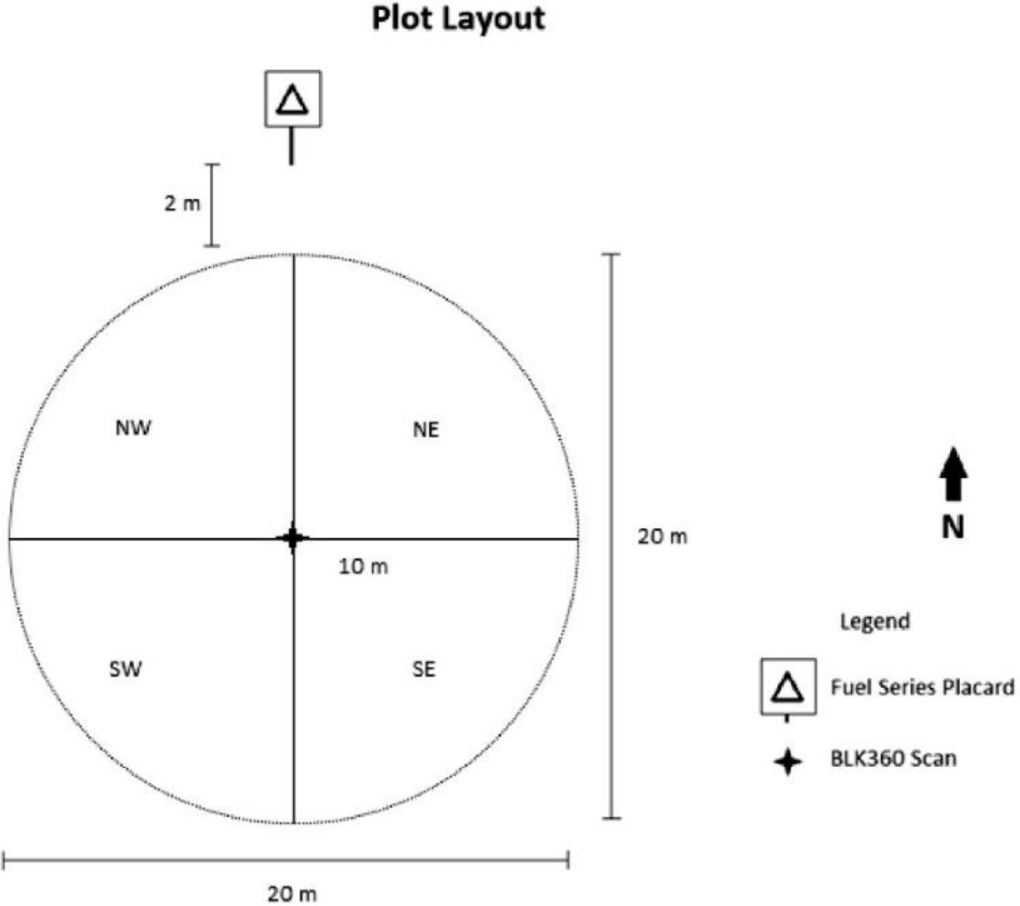
Plot layout for TLS monitoring field data collection methods. (Courtesy: Pokswinski and others 2021).

Once scanning is complete and the scanner is safely removed and stored, a prism pass is conducted using a 10-factor prism over plot center and targeted at 1.37-m (4.5 feet) above ground (at breast height). Trees are categorized as “in” or “out” and recorded as either conifer or broadleaved. The prism pass can eventually become optional for Tier 1 sampling after completing multiple iterations of Tier 2 sampling and site-specific models have been established. Periodical Tier 2 sampling, however, is optimal for crosschecking metrics and recommended after any major plot change (e.g. wind damage, thinning, etc.).

### Tier 2 Sampling - Transect & Overstory Data Collection

Tier 2 sampling includes both a Tier 1 scan and additional transect sampling. Once the TLS scanning is complete, fuel point intercept, coarse woody debris, fuelbed, litter and duff depth, and overstory data are collected, which requires establishing transects. The transects radiate 10-m out from the plot center in each cardinal direction, resulting in two perpendicular 20-m transects that intersect at plot center (Figure 4). Poles are placed at the terminal end of each transect with a tape measure tautly stretched along the transect. When establishing the transects, take care not to trample the vegetation or litter, because this material will be sampled. Depending on the complexity of the site and proficiency of the data collector, one can expect to collect the fuels and ecological conditions data in 30 minutes to 1 hour. See the IntELiMon site for example Tier 2 datasheets with fuel categories used in multiple ecosystem types.^2^

#### Fuel Sampling: Fuel Point Intercept Sampling

Point intercept sampling methods similar to those developed by Goodall (1952) can be used to estimate percentage of vegetation cover. An analogous approach is used here. Some fuel categories are standardized, such as live or fine fuels (graminoids, forbs, woody), leaf litter (conifer needles), and bare ground (no fuels) (Hiers and others 2007). Customizable categories can also be added, such as wiregrass, palmettos, bunch grasses, exotics, sagebrush, etc., Table 2).

**Table. 2.** Example point intercept fuel categories, as well as rules, & exceptions for a southeastern longleaf pine (*Pinus palustris*) Flatwoods ecosystem. This is an example of how fuel categories can be designed for a particular ecosystem type, using general (e.g. graminoids, woody live) and more custom (e.g., saw palmetto) fuel categories.

| **FUEL POINT INTERCEPT CATEGORIES** |  |
| --- | --- |
| **Graminoids** | Live and dead grasses, sedges |
| **Woody live** | Live woody trees and shrubs <10cm dbh |
| **Live saw palmetto\*** | Live saw palmetto (*Serenoa repens*) leaves and petioles |
| **Vines** | Live and dead climbing or trailing woody-stemmed plants |
| **Forbs** | Live and dead, detached and attached, small herbs that do not fit in any of the above categories |
| **Woody leaf litter** | Dead and detached non-conifer leafy material |
| **Pine needle leaf litter** | Dead and detached conifer needles |
| **Conifer litter** | Detached conifer material (bark flakes, catkins, etc.) |
| **Dead saw palmetto\*** | Dead saw palmetto (*Serenoa repens*) leaves and petioles |
| **Bare ground** | Only include dinner plate sized areas of bare ground, where no other fuel category is present or touching the fuel sampling rod. |
| * This is a custom category, created for the monitoring needs of the site and ecosystem type. |  |

| **FUEL POINT INTERCEPT SAMPLING RULES AND EXCEPTIONS** |
| --- |
| A fuel category can only be marked once per meter point intercept. For example, if an *Andropogon* and *Dichanthelium* grass both touch the rod on point 1.5m, only one tally would be marked for the graminoid group (even though both *Andropogon* and *Dichanthelium* are graminoids).<br>If the fuel sampling rod drops on a solid object (oak leaf, pinecone, stick, etc.) or covers up other vegetation, no other vegetation or fuels beneath this object are included in the tally.<br>Down woody debris (sticks, twigs, bark, stumps, logs etc.), pinecones, top-killed shrubs, trunks of trees, mosses, lichens are not recorded here. Down woody debris and pinecones, and trees (>10cm dbh) are recorded elsewhere.<br>If leaf litter is burned material, do not count as leaf litter if black and disintegrates when handled.<br>Bare ground is only tallied when soil at that point is not covered by any other vegetative material in a dinner-plate sized area or larger. If bare ground is tallied, then that is the only tally at that point. |

Starting 0.5-m from a terminal end and repeating every meter thereafter (0.5, 1.5, 2.5, etc.), the fuel sampling rod is vertically dropped along the tape measure. Vegetation material less than 1.4-m in height and touching the fuel sampling rod are tallied as present. Each fuel category is recorded once per sample point, such that if two grass species are present at the sample point the graminoid category will receive one tally. No other materials beneath an object are recorded if the fuel rod is not touching it. For instance, the rod may land on a large broadleaved leaf. Although there may be material below it (e.g. pine needles), they are not tallied unless they touch the rod elsewhere (above the broadleaved leaf). Bare ground represents no fuel and is only tallied when the fuel rod is touching bare mineral soil the size of a dinner plate that is not covered by any other vegetative material. When tallied, this should be the only tally at the meter point.

Upon completion, each transect will have 20 sample points for a total of 40 sample points in each plot. Each tally in a category will represent 2.5 percent cover for that category. The following are disregarded when using the sampling rod as they are accounted for elsewhere in the protocol: down woody debris (sticks, twigs, bark, stumps, logs), cones, and trunks of trees (see below). Mosses and lichens are not counted unless added as a custom category.

Below (Table 2) is an example of how point-intercept fuel categories can be designed for a particular ecosystem type, here a longleaf pine (*Pinus palustris*) Flatwoods ecosystem. Here, we used general (e.g. graminoids, woody live) and more custom (e.g., saw palmetto) fuel categories. In this example, “live saw palmetto” would only be marked under that category, not under “woody live.” And “woody live” accounts for all other live woody plants except saw palmetto. The idea is to start with the default fuel categories provided by the IntELiMon protocol and optionally expanded upon using customized fuel categories as desired or needed for monitoring needs.

#### Fuel Sampling: Fine and Coarse Woody Debris Sampling

Coarse woody debris sampling uses the planar intercept method (Brown 1974) to tally down and dead woody fuels that intersect the entire 40-m of transects. These fuels are broken out into standard fuel time lag categories (1-hour, 10-hour, 100-hour, and 1000-hour) and female conifer cones. A wildland fire fuel sizing gauge with a 6-mm slot for 1-hour fuels, a 2.5-cm slot for 10-hour fuels, and a 7.5-cm slot for 100-hour fuels can be used to determine the debris diameter. Along the two perpendicular 20-m transects a tally is made in the corresponding category each time one of these fuels crosses the plane of the transects. Note that one branch can be counted multiple times if it crosses the transect plane in multiple places. For example, one large branch with several small branches, several small individual branches, or the same large branch may cross the line more than once and tallied as such. For this sampling, decay status is disregarded and does not affect count or diameter. Standing dead material attached to the ground (i.e., top-killed shrubs) is tallied if it is less than 1.5-m in height. Each individual female conifer cone that crosses the transect plane is tallied and the species is noted, if possible.

#### Fuel Sampling: Fuelbed, Litter, and Duff Depth Sampling

Fuelbed, litter, and duff depths are collected at 0.5, 3.5, 6.5, 13.5, 16.5, and 19.5-m along each transect for a total of 12 measurements. The fuelbed depth is defined as the top of the litter layer (Oi) to the tallest live or dead plant material. The fuelbed depth is sampled across a plane 30.48-cm (12 inches) on each side of the measurement point perpendicular to the transect tape, with a maximum fuelbed depth of 1.4-m in height. The maximum height on each side is used to estimate an average fuelbed height along that perpendicular plane and recorded at that point. At this same sample point, the litter depth is recorded as the greatest height where litter touches the rod. Note that ‘litter’ can either be defined as ‘leaf litter’, ‘fine litter’, or ‘dead fine vegetation’ depending on the ecosystem type and monitoring needs. The duff layer depth, containing partially to highly decomposed material, is also measured ending at the top of the mineral soil. Care must be taken to avoid compressing the material when measuring this layer. Measurement increments are 0.0, 0.1-cm (for trace organic material), 1.0-cm, and by the centimeter thereafter.

#### Overstory Sampling

Overstory trees larger than 10-cm in diameter at breast height (d.b.h.) are characterized within the four quadrants 10-m of plot center (delineated by perpendicular transects). Trees are tallied by quadrant (NW, NE, SE, and SW), species/taxa, and quantity.

### Tier 3 Sampling - Additional Data Collection

Tier 3 sampling is performed as needed, or with specific objectives in mind; for instance, to measure surface fuel mass in clip plots (e.g., Loudermilk and others 2023), burn severity (e.g., Gallagher and others 2021), or woodland/rangeland woody cover via line intercept methods (e.g., Herrick and others 2005) to relate to the TLS output metrics. Other examples include additional tree measurements, biodiversity sampling, destructive harvesting, or qualitative information such as habitat quality.

### Data Offloading

At the end of each collection day, export scans and delete them from the BLK360’s internal storage. Using a fully charged battery can prevent the BLK360 G1 from shutting off during the data exporting process. Refer to the latest Methods Manual for step-by-step instructions for using BLK Data Manager Utility and Cyclone Register to export, delete, and convert scans from the proprietary ‘.blk’ format to more widely usable ‘.ptx’ format.^2^

## OUTPUTS AND METRICS

### Automated Processing and Analysis of Data

The TLS data collected through this protocol can be processed and analyzed automatically through the Interagency Ecosystem LiDAR Monitoring (IntELiMon) viewer, a streamlined processing pipeline maintained by the USGS Earth Resources Observation and Science (EROS) Center (USGS 2023) (Figure 5).^3^ TLS scans can be directly uploaded to IntELiMon, on the EROS server, where they are processed through carefully developed scripts and archived with explicit metadata for each dataset. Scan file and plot ID preparations are described in the Methods Manual.^2^ This initial data flow generates hundreds of structural metrics calculated from each TLS 3D point cloud and, when coupled with initial field transect data (from Tier 2 sampling), is used in a machine learning environment to predict tree and fuels data for forestry and ecological monitoring. EROS is capable of processing 100 TLS scans per day and results can be accessed via the IntELiMon Viewer, an interactive data dashboard for users to access and download processed TLS data and metadata. Quality control/quality assurance and metadata management are built into the data workflow.

**Figure 5.**
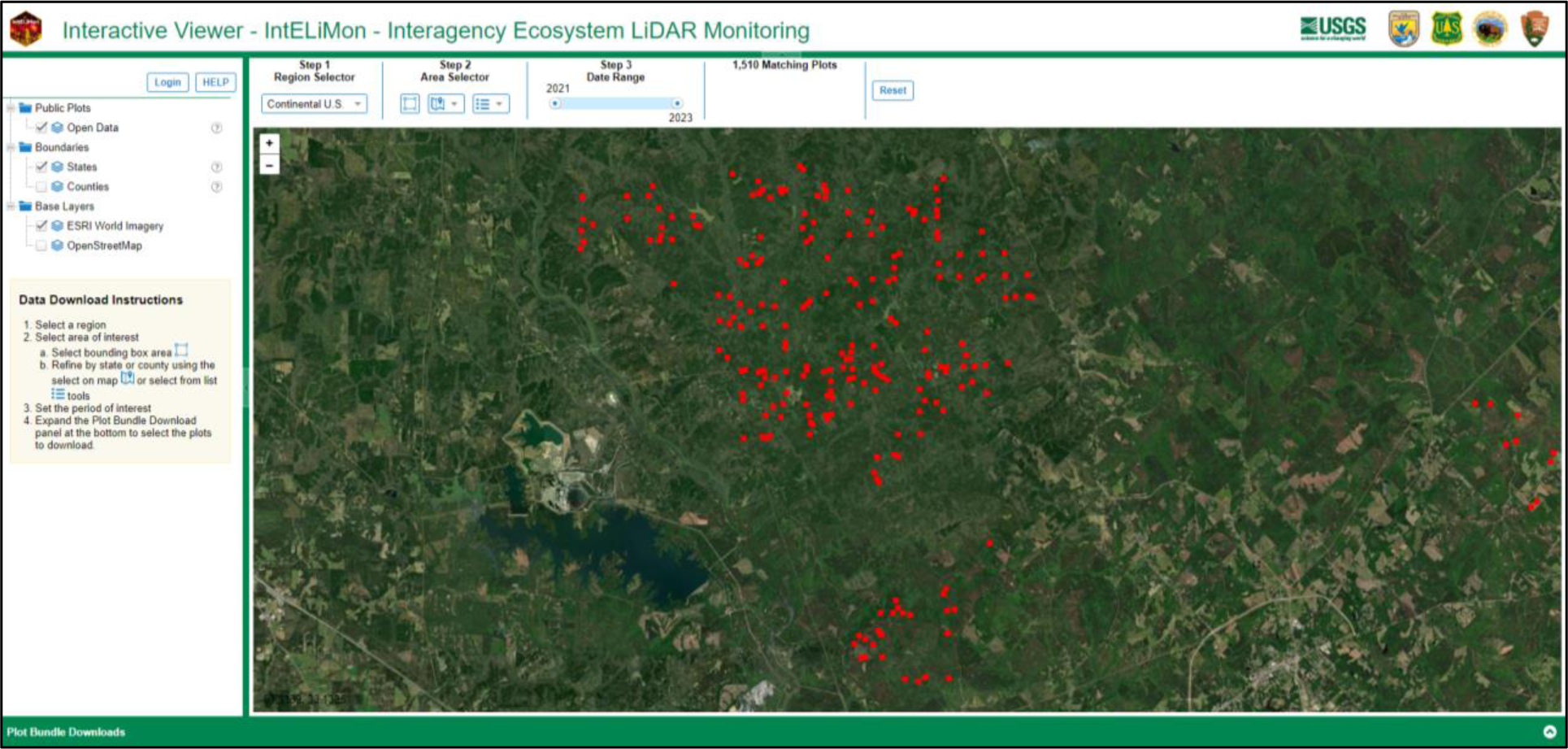
IntELiMon Interactive Viewer dashboard (as of March 2024) (USGS 2023).

Metrics derived from TLS scans capture the complex ecological structure of a plot. Given the millions of data points within each point cloud, multiple ways exist to measure and characterize portions of the scan. When processed with the EROS scripts, scans are clipped to a 15-m radius, normalized (elevation measurements are converted to height above ground), and, depending on the metric, are analyzed in their entirety, within designated height strata, or within 2-cm by 2-cm by 2-cm voxels (a 3D pixel) or cubic space. Surface vegetation is defined in stratums of 0.5, 1, 1.5, 2, and >2 m, and forest stratum types include groundcover (0–1 m), understory (1–3 m), midstory (3–9 m), and overstory (>9 m). Using a spherical neighborhood, the point cloud is rasterized and analyzed for summary metrics, including the proportion of returned and unreturned pulses and occluded space. Here, the x,y,z coordinates are converted to spherical coordinates. This allows points to be tracked along their flight path and for estimation of the proportion of pulses that return to the scanner from a particular distance, those that continue further away from the scanner, and those that never return (i.e. never hit an object and return). This method can be used to differentiate between true gaps (i.e., space without vegetation) and occluded space (i.e., not measured because no pulses passed through the volume). The input .ptx point cloud is also converted to .las format for basic LiDAR outputs, including a DTM, CHM, summary statistics, and height bin metrics. A tree classification identifies individual boles via a segmentation algorithm (de Conto and others 2017), providing a tree inventory and metrics that include corrected basal area, tree descriptive statistics, and foliage diversity metrics. Fuels are filtered out to create a stemless 0–3-m point cloud, allowing for shrub classification and output shrub inventory and summary statistics. Fine fuels and coarse woody debris are further defined from the fuels point cloud. This results in over 200 forestry metrics and structural characteristics, tree and shrub inventories, and raster grids, including a DTM, CHM, and 10-m^2^ shrub tile (Table 3, Figures 6, 7, and 8). These metrics range from vegetation distribution, density, tree and fuel heights, and d.b.h. to forest structural characteristics including differences between true forest gaps and space created by occlusion. Note that metric improvement and development is continuous. As metrics are updated, previously collected TLS scans and field data can be reanalyzed for metric and data updates and consistency. Details on processing and early metric development are found in Gallagher and others (2021), Maxwell and others (2023), and Loudermilk and others (2023).

**Table 3.** Examples of what is directly measured (using Tier 1) or predicted (using either Tier 2 or Tier 3*) metrics from the TLS monitoring protocol. TBD means that the measurements or estimates could still be developed. N/A means that it is not applicable. The ’New category’ represents the ability to add new categories/metrics/predictors as they are developed. d.b.h.: diameter at breast height, c.b.h.: canopy base height. The ‘Other Nuanced Categories’ represent the multitude of categories that are not necessarily directed related to a known vegetative feature but quantify various portions of the point cloud using statistical measures. These measures quantify the structure of the vegetation, space between vegetation, and occlusion behind vegetation.

| CATEGORY | DIRECT MEASUREMENTS | PREDICTED/CALIBRATED MEASUREMENTS |
| --- | --- | --- |
| **Trees** | Tree density, % canopy cover, height, d.b.h., c.b.h. | Conifer vs. Broadleaved |
| **Shrubs** | Shrub density, heights | % cover |
| **Graminoids** | TBD | % cover |
| **Leaf debris** | TBD | % cover |
| **Woody debris** | Estimates of 1,10,100,1000 fuels | % cover |
| **Duff** | TBD | Depth |
| **Fuelbed** | < 3m height estimates | Depth |
| **Surface fuel loading (<1m height)** | N/A | Surface fuel biomass (tons per acre)* |
| **Vegetation/fuel volume** | Volume of canopy, ladder, & surface fuels | Volume of canopy, ladder, & surface fuels |
| **New category (as developed)** | TBD | TBD |
| **OTHER NUANCED CATEGORIES** |  |  |
| **Height statistics** | Height estimates in various portions of the TLS point cloud | TBD |
| **Space & occlusion statistics** | % of space in various portions of the TLS point cloud | TBD |

**Figure 6.**
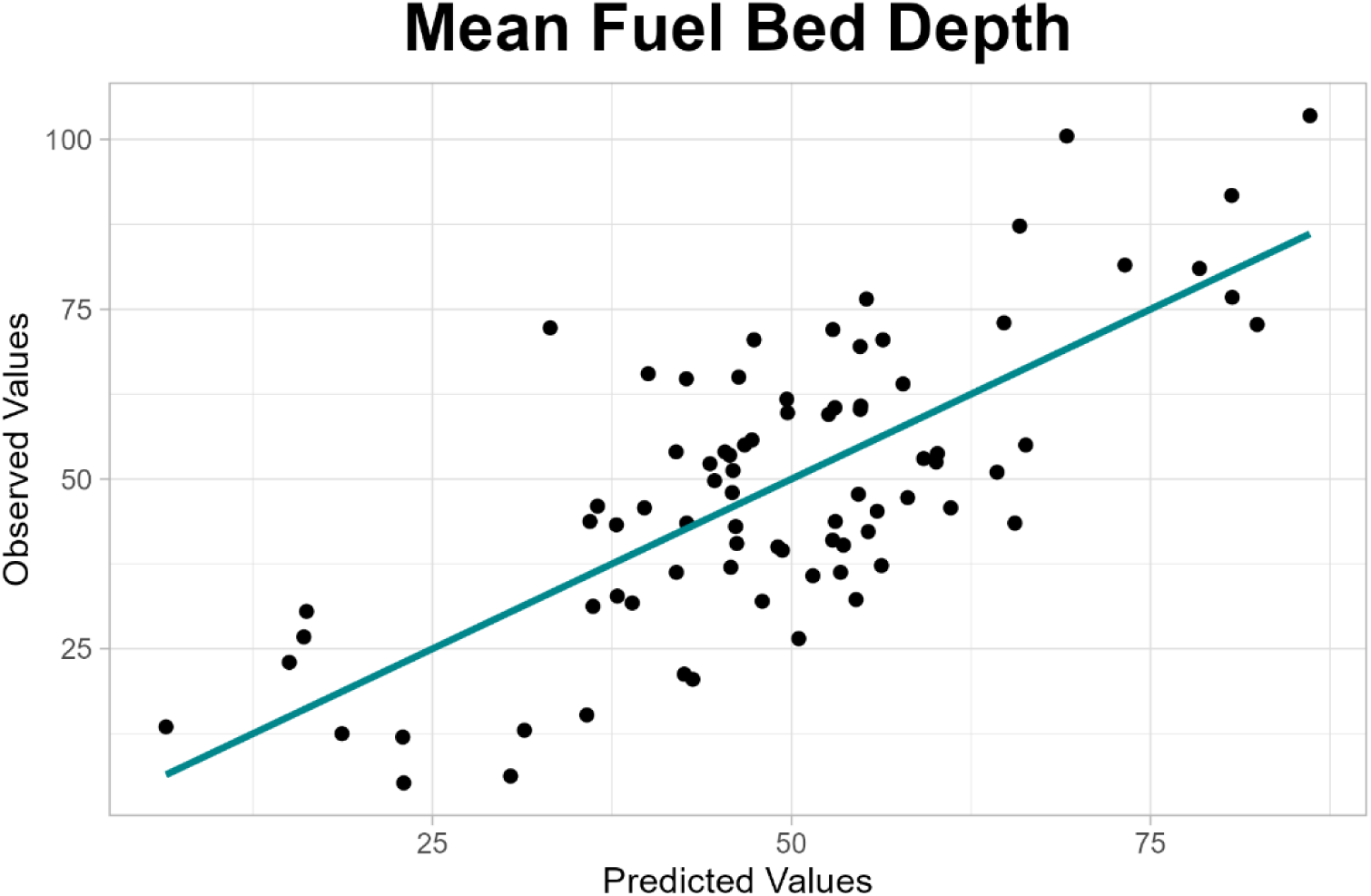
Mean fuelbed depth as predicted using the TLS metrics (from Tier 1) to predict field sampled plot-level mean fuelbed depth (from Tier 2) using linear regression. Units are in centimeters.

**Figure 7.**
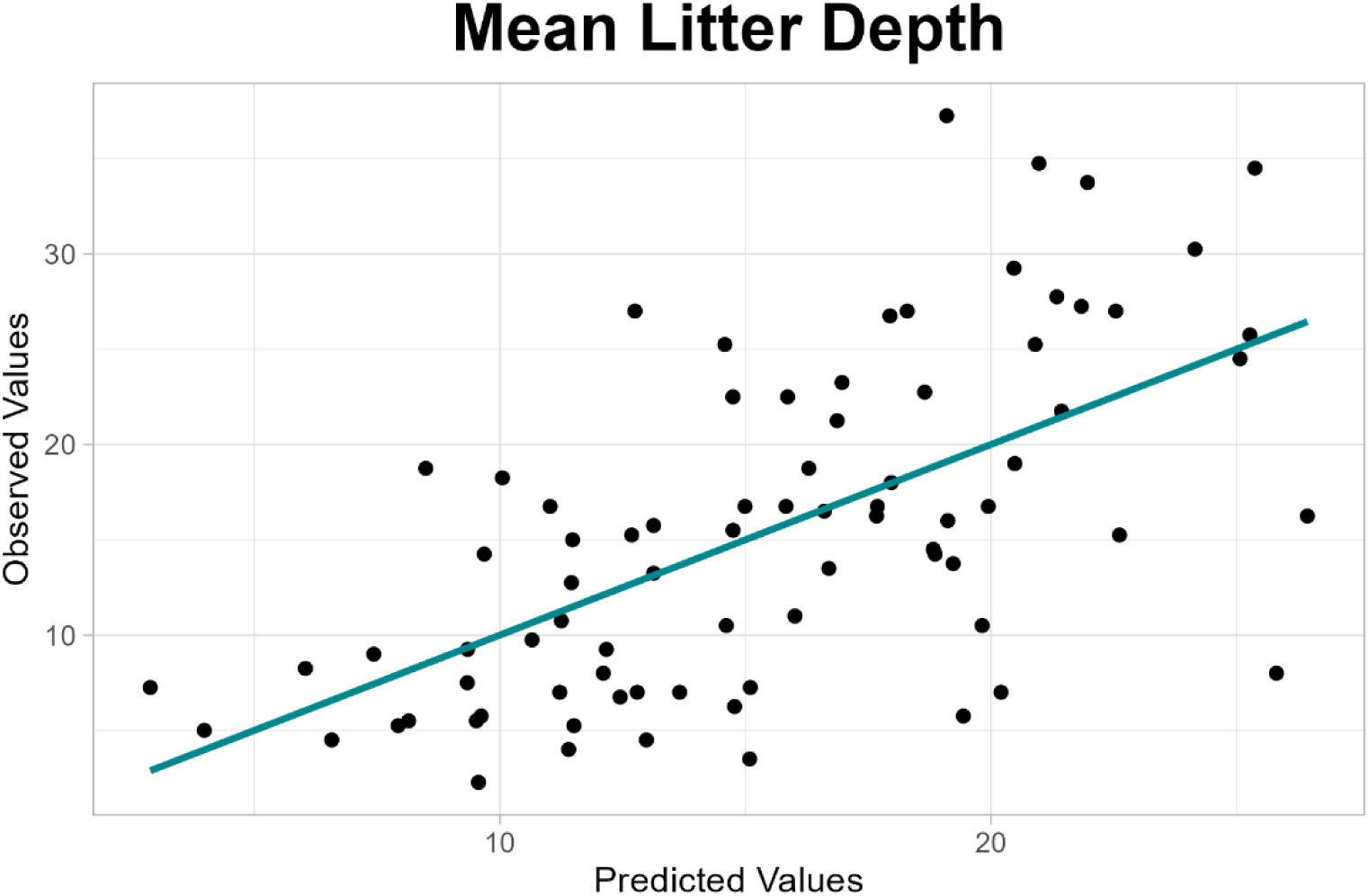
Mean litter depth linear as predicted using the TLS metrics (from Tier 1) to predict field sampled plot-level mean litter depth (from Tier 2) using linear regression. Units are in centimeters.

**Figure 8.**
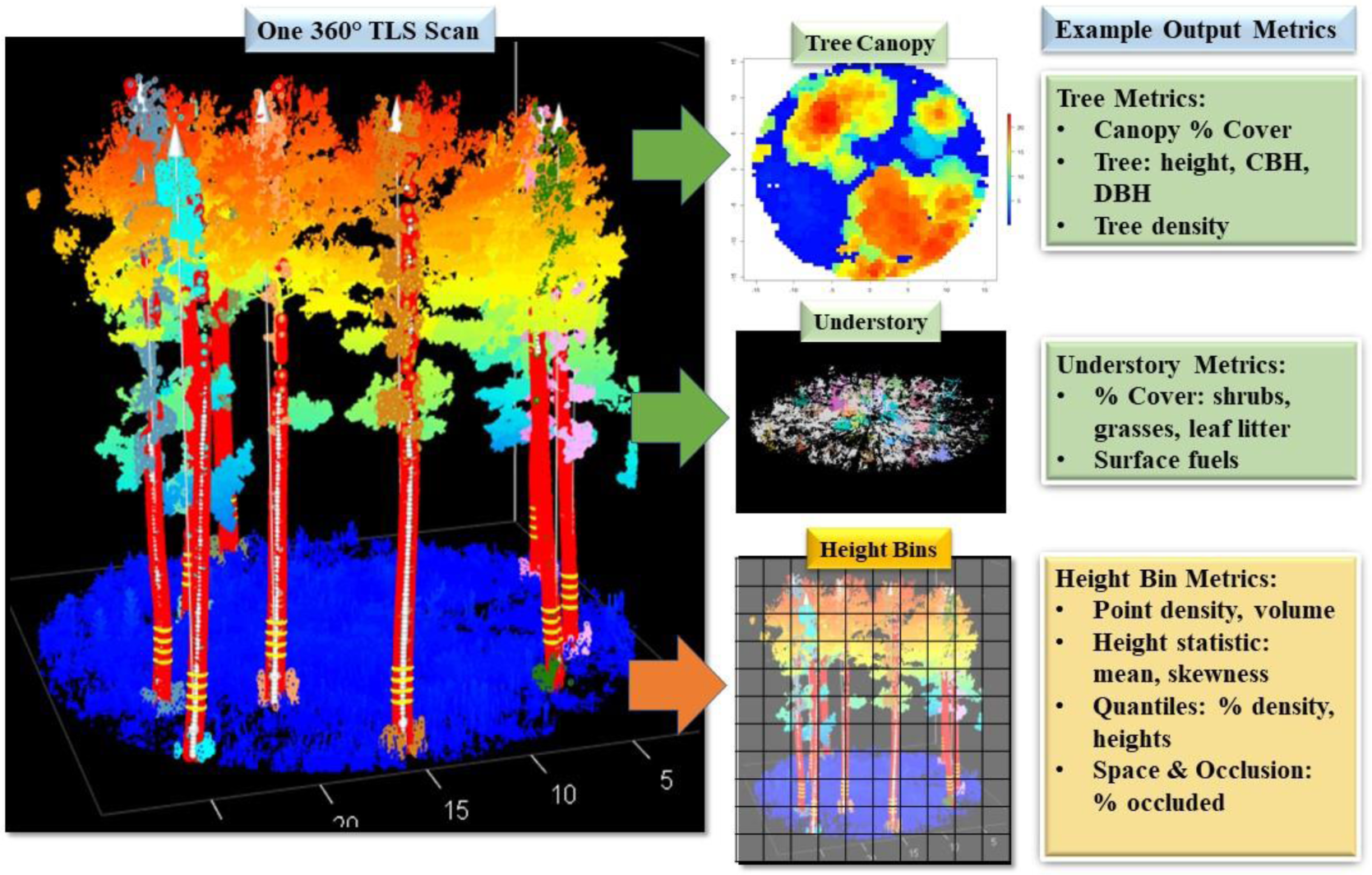
Processing flow: Example of one 360° terrestrial laser scan as output by the IntELiMon system. Trees are located, identified, and measured for diameter, height, and overall canopy cover, then removed digitally to estimate understory or percentage of surface cover vegetation and fuels (green boxes). Then the entire 3D dataset (point cloud) is segmented into height bins (or 3D boxes) and hundreds of metrics are calculated (yellow boxes) to use in predictive modeling (e.g., surface biomass, burn severity). Furthermore, canopy, ladder, and surface fuel volumes are calculated. Please also refer to Table. 3.

### TLS-Based Monitoring Challenges

The most notable limitation to TLS-based monitoring is the obscuration caused by objects, such as large trees or dense vegetation close to the scanner. This “occlusion” blocks the laser pulse from collecting data behind a given object. Occlusion is accounted for in this monitoring protocol using several methods. First, occlusion is calculated from the point cloud, where all laser pulses (regardless of whether they intercept an object or not) are accounted for. This allows for the differentiation between true empty space (e.g., open canopy space) versus obscured space (e.g., space behind a tree bole). This also applies to the differentiation between true and unknown vegetation structures. Second, there is ongoing research to predict what is being occluded given the existing plot structure captured by the scan. Although there are well-known approaches to merge or “stitch” multiple adjacent scans together to provide a complete 3D point cloud of all the vegetation [for examples, see Cheng and others (2018), Guan and others (2020), Tremblay and Beland (2018)], the loss of efficiency needed to merge multiple scans currently outweighs the potential gain in information. Merging multiple scans is further complicated by vegetation movement during and between scans, causing errors in the merging process and output metrics. The dense structural information in a single scan, even given obstruction issues, ultimately provides millions of structural 3D data points to work from. This point-based monitoring approach is more straightforward than attempting to merge multiple scans, increasing efficiency of data collection and processing, notwithstanding standardization of sampling. There are however ongoing research studies to streamline this merging process.

Ground-level data, including compact leaf litter, organic soil layer (duff), or microtopography are not currently as well characterized as larger objects (trees, shrubs) in the scans. The Tier 2 sampling data (e.g., transect sampling) are used to overcome this bottleneck by building site-level models to predict these lower strata data from field data as calibration. Fine or coarse woody debris can be captured given the high point density of the scans and distinct structural patterns of the wood, but is dependent on location within the scan, vegetation density, and size of the debris. Again, the Tier 2 sampling data are well suited to build site-level predictive models of this information. Standing dead tree or snag identification is currently a limitation, although they can be differentiated in leaf-on or strictly evergreen conditions. Leaf-on versus leaf-off scans of the same plots also provide information on snags.

Currently, the Leica BLK360 series is supported for this TLS monitoring protocol, however, there is already promise in using other scanners (Stovall and Atkins 2021). As technology advances, new instruments will be tested.

## RECENT RESEARCH AND APPLICATIONS

### Recent Research

Extensive research has contributed to the development of this TLS monitoring protocol, including metric analyses and methodologies blending TLS and field sampled data. Here we review the most recent publications that use the BLK360 G1 scanner for measuring complex structural characteristics (metrics) of terrestrial ecosystems at the plot level and further prediction modeling of other structural and nonstructural aspects of the system. See references throughout this section for further details on methodologies and analyses not covered in this paper. These advances drive the operational adoption of TLS for long-term monitoring and management by continuously improving point cloud metric measurement and interpretation. It should be noted that these are the most recent examples to date using BLK360 G1 scanners and that technology and methods are continuously advancing (see Future Directions of Research for examples). Many other TLS and LiDAR applications in ecosystem and forestry research are not included here as they are beyond the scope of this paper.

BLK360 G1-derived metrics were initially developed by Anderson and others (2021) and Gallagher and others (2021). Both collected field measurements to relate to the metrics calculated from each individual scan using a BLK360 G1. The metrics were similar in measuring trees, shrub, and herbaceous cover, as well as calculating density and height statistics within height classes. They differed in how they divided the scan point clouds into height bins and calculated percentage of various types of vegetation cover. Anderson and others (2021) used a combination of TLS and traditional field-based sampling methods to predict groundcover species richness in southern pine Flatwoods. Vegetation structure, assessed as vertical and horizontal openness and variability, was used as a proxy for richness in the TLS data in addition to intensity, or the strength of each return. TLS metrics were found to accurately predict species richness in groundcover plant communities where structure and richness are tightly coupled. They found that herbaceous richness was generally more accurate than shrub richness. Gallagher and others (2021) used the single-scan TLS approach for monitoring wildland fire burn severity in the New Jersey Pinelands National Reserve. They found that TLS-based methods for quantifying plot-level burn severity offered significant improvements compared to the Composite Burn Index (CBI), the most widely used field-based method. The authors found that TLS overcame CBI’s limitations of subjectivity and repeatability and excelled at characterizing understory vegetation where prescribed fire effects are most pronounced, where both CBI and ALS are weakest due to qualitative estimates and canopy occlusion, respectively. That same year, Pokswinski and others (2021) published the first methods paper using the BLK360 G1for terrestrial ecosystem monitoring and introduced the initial plot-based, transect methods featured in this paper.

Loudermilk and others (2023) demonstrated TLS capabilities for predicting understory vegetation and fuel biomass as well as consumption by fire in frequently burned Southeastern United States’ pinelands. They describe 162 of the TLS metrics in detail, and illustrate how various fuel mass classes, such as fine fuels, fine woody debris, or simply total surface biomass pre- and post-burn, can be predicted using these TLS metrics. Finally, Maxwell and others (2023) assessed shrub height predictions from TLS metrics (in the New Jersey Pinelands National Reserve) when (1) varying the number of randomly selected shrub height measurements and (2) using metrics from leaf-off versus leaf-on TLS data. They found that at least 10 random shrub heights (20 being ideal) were needed for accurate predictions and that leaf-on TLS data generally provided more precise predictions. Their work highlights the importance of reference data sampling density, design, and data characteristics when training empirical models for extrapolating to new sites or plots.

Other notable references are related to single or multi-scan analyses using other TLS instruments. Batchelor and others (2023) illustrate the use of single scans of the FARO® Focus 3D S120 terrestrial laser scanner (FARO Technologies Inc., Lake Mary, FL, USA) for estimating new structural complexity metrics in Pacific Northwest forests. Many studies have used multiple merged scans to predict grass, shrub, tree, or fuel attributes at various scales and using a variety of metrics of interest (e.g., Alonso-Rego and others 2020, Cooper and others 2017, Loudermilk and others 2009, Olsoy and others 2014, Wallace and others 2022). Merged FARO® Focus scans were used to relate structural complexity of mixed pine-hardwoods forests of the southern Appalachians to vascular plant biodiversity (Walter and others 2021). Stovall and Atkins (2021) compared merged predictions of tree metrics output and modeled from the BLK360 G1 and the FARO® Focus 3D S120. They concluded that while metrics were comparable between instruments, laser technology (BLK360 G1: time-of-flight versus FARO® Focus: phase-shift) impacts each scanner’s range and point noise output. Generally, the BLK360 G1 had less noise and required less filtering than the FARO® Focus.

### Future Directions of Research

Research informing this monitoring protocol continues to advance with the development of new methods and applications for 3D forest structure characterization. LiDAR technology is also advancing quickly and instruments well-suited for this type of monitoring continue to be developed. Therefore, understanding how these instruments sample the environment will be vital for TLS-based monitoring to maintain consistency and value in the future. Future research in this area will include direct comparisons of new and old sensors to set a standard that can be maintained. This includes comparing the precision of the BLK360 G1 scanner with other low-cost alternatives (Stovall and Atkins 2021). Such comparisons can assist in addressing current limitations of TLS monitoring and tailor scanner instruments for unique ecosystems and monitoring needs (See TLS-Based Monitoring Challenges).

Metrics derived from the 3D point cloud are continually improving in detail and accuracy. This includes predicting occluded areas given the existing plot structure captured by the scan. There is ongoing research on segmenting point clouds into different types of fuels (e.g., coarse woody debris versus leaf litter) or mapping shorter vegetation, such as tree seedlings, shrubs, or bunchgrasses with promising results. There is also an effort to create 3D tree, shrub, and grass models in an inventory or 3D library to generate virtual vegetation or forested stands to simulate and quantify a desired or representative fuel environment. Quantitative structure models (QSMs) are particularly useful for such segmentation work (Brede and others 2019, Stovall and Atkins 2021). QSMs and other advancements can distinguish between plant species or plant functional types based on their relative structural characteristics (e.g., broadleaved versus needle-leaved trees). QSM approaches can also provide unique insights related to species identification (Åkerblom and others 2017, Krooks and others 2014) and plant physiology and structure (Hackenberg and Bontemps 2022). There is a need for understanding fine-scale complexities, such as further characterization or prediction of groundcover plant diversity or soil organic layer development and consumption by fire. Other structural characteristics captured by the 3D point cloud, such as canopy density, canopy gaps, ground substrate, and soil surface characterization have additional applications, including erosion models.

Future integration of TLS with ALS and spectral reflectance datasets could provide full vegetation distribution datasets for more complete landscape analyses. Work is being done to automate the processing of multiple overlapping TLS scans and overlapping TLS and ALS scans to expand the coverage of TLS fine-scale data to the extent of ALS through imputation and interpolation techniques. There is potential to integrate a spectral-reflectance index with the 3D data for even further improvements to fine-scale burn severity mapping as well as identifying species, plant functional types, or fuel types. Another major research avenue is to apply TLS vegetation and fuels data, including 3D libraries, as onramps for next generation wildland fire and smoke simulation models such as WFDSS, FIRETEC, and QUIC-Fire (Linn and others 2020, Loudermilk and others 2023, Peterson and others 2022). By providing detailed wildland fuel parameters and forest structure, particularly in 3D, TLS monitoring data can be used as a direct input, or coupled with ALS, for full coverage of canopy and understory vegetation characterization. As such, 3D point clouds can be used to develop 3D fuels inputs to advance fire models (Mueller and others 2021). The high detail from TLS is uniquely suited to laboratory scale experiments and comparisons with such models, where it can identify thresholds in fuel mapping detail and help inform fine scale fire spread processes (Marcozzi 2022).

### Management Applications and Demonstration

Numerous land management agencies, including the Forest Service, U.S. Fish and Wildlife Service, National Park Service, and Bureau of Indian Affairs, amongst others, currently implement TLS monitoring, with other groups expressing interest in its use. The FWS Southeast Fire Program first unveiled this protocol for fire effects monitoring of hazardous fuel treatments on national wildlife refuges in collaboration with the Forest Service and university research partners. Although the effort began in the Southeastern United States, it is quickly gaining national interest, and operational training and testing has occurred across additional national wildlife refuges, national forests, national parks, tribal lands, and Department of Defense installations, amongst others. More training events are planned at national training centers (See Training, Coproduction, Technology Transfer).

The protocol arose out of a need to improve the accuracy and precision of fire effects monitoring and is thus well-suited for capturing changes in understory vegetation structure and volume due to both natural disturbances and management treatments. Pre- and post-fire TLS monitoring can be used for surface biomass change detection to estimate fuel consumption and assess maintenance of target surface fuel characteristics over time (Loudermilk and others 2023). Other demonstrated applications of the TLS protocol include critical habitat evaluation, documentation of restoration projects or disturbance events, and for determining habitat suitability for potential use or wildlife translocation. For example, on St. Marks National Wildlife Refuge, TLS monitoring is used to assess habitat of threatened and endangered species such as the frosted flatwoods salamander (*Ambystoma cingulatum*), black rail (*Laterallus jamaicensis*), and red-cockaded woodpecke*r (Dryobates borealis*) and to quantify response efforts to bark beetle disturbances (Link 2023). Other applications include, but are not limited to, assessing changes through time, such as forest succession, mortality or regeneration patterns, fire effects including fuel consumption, invasive species encroachment, or management treatment response.

In addition to its monitoring applications, TLS has significant potential to inform next generation national fuels maps, benefiting fire models and decision-support systems that rely on such data. Current national fuels maps such as LANDFIRE and decision-support systems such as WFDSS primarily use data derived from 30-m satellite imagery but are limited in their ability to use detailed 3D fuels data. In contrast, new prototype fuel mapping systems such as FastFuels (USDA Forest Service 2023), fire models such as QUIC-Fire, and decision-support systems that integrate them, such as the BurnPro3D platform developed by WIFIRE (UC-San Diego 2023), all represent fuels in high detail 3D, and can thus leverage 3D data not only from TLS but from ALS or UAS as well. Each of these 3D mapping technologies has its own strengths and limitations and next-generation fuels maps could likely integrate all of them. Although TLS is not wall-to-wall in coverage, its below-canopy detail will greatly enhance fine-scale surface fuel characterization. Integration of detailed 3D fuels data and fire models could lead to a wide range of management applications outside of current capabilities, including simulation-based assessments of fuel treatments, testing prescribed burn ignition patterns, and evaluating risk in the context of prescribed burns. Natural resource managers in many landscapes are now demonstrating this technology for use in monitoring and working with scientists and practitioners for advanced innovation and research. Ultimately, these new technologies could serve to advance decision-support tools needed for prescribed fire or fuel treatment planning, silviculture prescription design, assessments of fire risk and fire effects, and other natural resource management needs.

Beyond the valuable ecological metrics provided by TLS monitoring, RGB panoramic photos captured by the BLK360 G1 scanner serve as an incredible visual communication tool. They can be collected from the scanner alongside the point cloud and give a detailed look at a given plot from a human perspective. Flat images are notably lighter weight than point clouds, consuming at least two orders of magnitude less disk space per file. As such, panoramic imagery is a more viable option for sharing between colleagues, especially from the field over a cellular network. WIFIRE, a University of California-San Diego-based program specializing in data integration and visualization of wildland fire behavior, has created immersive visualizations of TLS monitoring plots by reprojecting the TLS point clouds to an RGB panoramic image. These “immersive forest experiences” allow the user to virtually revisit monitoring plots, witness the effects of management treatments, and easily recollect structural attributes. In addition to displaying the RGB colors collected from the BLK360 G1, the ecological metrics can be embedded and displayed in the immersive experience, giving the user the opportunity to see their metadata in context.

### Training, Co-Production, & Technology Transfer

Training, coproduction, and technology transfer opportunities for utilizing this TLS monitoring protocol are well underway across the United States. While training is currently a collaborative grassroots effort among Federal agencies and research consortia, more formalized training curricula are in development. Check with each agency’s training or technology transfer liaison for upcoming opportunities. Courses range from technical training on field data collection and initial data processing (see Field Data Collection Methods and Outputs and Metrics) to data analysis and modeling for advancing the application of TLS monitoring in land management decision support.

To accelerate technology transfer for innovative land management tools, the Department of Defense and USGS, in partnership with a host of other Federal partners including the Forest Service, have established the National Innovation Landscape Network (NILN)^4^. Currently, the Innovation Landscape Networks formed in both the Southwestern and Eastern United States (EILN)^5^ have made TLS monitoring and fuels characterization a focal area for innovation and technology transfer. Co-production is recognized as the key to success for this type of innovation, where technology and management needs hit a crossroad, but can efficiently and effectively advance with continuous feedback and involvement between researchers and resource managers. As these innovation landscapes develop and mature, opportunities for scaling up training, technology transfer, and novel uses for TLS will likely grow and expand over time.

## Acknowledgements

This work was a collaborative effort developed with partners from the U.S. Department of Agriculture: Forest Service, U.S. Department of Interior: U.S. Fish and Wildlife Service, National Park Service, Bureau of Indian Affairs, and U.S. Geological Survey, U.S. Department of Energy: Los Alamos National Laboratory, New Mexico Consortium, West Virginia University, Northern Arizona University, and University of California – San Diego. We thank the Department of Defense’s Strategic Environmental Research and Development Program and the Environmental Security Technology Certification Program for their continued support. Seth M. Munson and Kurtis Nelson of the U.S. Geological Survey contributed to the development of this monitoring protocol and LiDAR analysis methods. The use of trade or firm names in this publication is for reader information and does not imply endorsement by the U.S. Department of Agriculture of any product or service. IntELiMon is an ever-evolving monitoring protocol and website where processing, field protocols, instrumentation, and data products are continuously updated.

## Footnotes

1 https://dmsdata.cr.usgs.gov/lidar-monitoring/

2 https://dmsdata.cr.usgs.gov/lidar-monitoring/monitoring

3 https://dmsdata.cr.usgs.gov/lidar-monitoring/viewer/

4 https://serdp-estcp.mil/page/38f0be40-b397-446f-bf1f-a401fe12423f/national-innovation-landscapes-network

5 https://www.fs.usda.gov/research/srs/projects/eiln

